# Stroke-Related Changes in Tonic and Phasic Muscle Recruitment During Reaching Reveal Pathway-Specific Motor Deficits

**DOI:** 10.1101/2025.05.28.656732

**Authors:** Anna S. Korol, Amelia Adcock, Valeriya Gritsenko

## Abstract

Upper limb motor deficits are common after stroke and often persist despite rehabilitation. While clinical assessments emphasize movement quality, they do not capture the underlying neuromuscular impairments, particularly in individuals with mild deficits. This study aimed to characterize stroke-related changes in muscle recruitment during reaching by separating tonic (gravity-compensating) and phasic (intersegmental dynamics-related) components of EMG activity.

We recorded surface EMG from 12 upper limb muscles during goal-directed reaching in 8 individuals with unilateral ischemic stroke and 9 controls. Using principal component analysis, we extracted tonic and phasic components and compared their amplitude, directional tuning, and coactivation patterns across groups. Group differences were evaluated with generalized linear mixed-effects models, regression, and correlation analyses.

Even individuals with mild stroke exhibited abnormal muscle recruitment. Proximal muscles were over-recruited in directions that typically require less activation, indicating altered directional tuning. Phasic activation of distal muscles was significantly reduced and worsened with time post-stroke (R^2^ = 0.52, p = 0.002). Tonic overactivation of proximal muscles was present across all stroke participants. Muscle coactivation patterns were hemisphere-specific: right-hemisphere stroke reduced tonic coactivation in contralateral arms, whereas left-hemisphere stroke increased it. Abnormal phasic coactivation between proximal and distal muscles correlated with impaired intersegmental dynamics compensation (R^2^ = 0.67, p = 0.013). Tonic and phasic impairments were often correlated, suggesting shared disruption of corticospinal and reticulospinal pathways.

These findings reveal distinct yet interacting deficits in muscle recruitment following stroke, supporting the development of neuromechanically-informed tools for individualized rehabilitation.

## Introduction

Stroke is the third leading cause of combined mortality and disability worldwide ^1^, with upper limb motor impairment affecting 80% of survivors ^2^ only half of whom regain useful upper limb function.^3^ Despite advances in rehabilitation, many individuals continue to experience poor arm control even years after stroke onset. Clinical assessments often focus on observable movement quality or joint angles, which cannot assess fatigue and may overlook the underlying neuromuscular coordination strategies that shape recovery.^4^ Understanding how stroke alters the recruitment of specific muscle groups, especially in response to biomechanical demands such as antigravity support and intersegmental dynamics, is essential for developing more targeted individualized rehabilitation strategies.^5,6^

Clinical assessments rely on low-resolution scoring and rater training to reduce variability, but this limits responsiveness and predictive validity—especially in patients with mild deficits—due to ceiling effects.^7,8^ These tools also cannot distinguish impairments from passive mechanics versus active muscle coordination, leading to inconsistent treatment outcomes in trials.^9–11^ We previously showed that motion capture– based measures of muscle torque are more sensitive than joint angles in detecting post-stroke motor deficits.^4^ This is especially valuable for patients with minimal visible impairment who still report difficulty. The current study builds on this by analyzing muscle recruitment during goal-directed reaching, aiming to identify objective physiological markers that improve on current clinical assessments.

Abnormal muscle recruitment patterns after stroke lead to inappropriate force generation that underlie motor impairment. During reaching, muscle forces serve two primary functions: (1) supporting the arm against gravity—primarily mediated by proximal muscles that span the shoulder and elbow, and (2) compensating for complex limb dynamics—including inertial, Coriolis, and centrifugal forces—primarily mediated by distal muscles that span the elbow and wrist. Muscle forces also generate stiffness and viscosity that help stabilize joints and respond to unexpected perturbations—features that are not fully captured by muscle torques alone. Muscle forces result from muscle contractions caused by motoneuron firing, which is observable with EMG. In our previous work, we demonstrated that the tonic component of EMG reflects muscle forces related to gravity compensation during reaching, while the phasic component reflects the remainder forces related to intersegmental dynamics.^12^ After stroke, efforts to actively support the arm against gravity or to load the shoulder often result in coupled elbow moments, which reduce the reachable workspace and restrict movement range.^13–16^ This pattern suggests that neural mechanisms for gravity compensation—likely mediated by the reticulospinal system—may become abnormally engaged after stroke.^17–19^ In addition, stroke disrupts complex intralimb coordination and grasping^6,14,20^, functions that may depend on plasticity within corticospinal projections.^21–23^ The reticulospinal and corticospinal systems are hierarchically organized and anatomically coupled,^24–26^ further complicating efforts to develop targeted interventions. In this cross-sectional feasibility study, we investigate how stroke affects these systems by testing two hypotheses: 1) stroke affects the recruitment of distal muscles that move the wrist and hand more than proximal and biarticular muscles that move the shoulder and elbow; 2) stroke affects the phasic component of muscle recruitment responsible for intersegmental dynamics compensation more than the tonic component of muscle recruitment responsible for gravity. Testing these hypotheses will result in clinically meaningful insights into post-stroke recovery and provide a foundation for individualized motor rehabilitation approaches.

## Methods

### Data Collection

Participants were grouped as Control and Stroke. The Control group included nine healthy, right-hand dominant individuals (mean age ± SD = 22.78 ± 0.67 years; 3 females, 6 males). The Stroke group included eight individuals (mean age ± SD = 58 ± 6.9 years; 1 female, 7 males) with single unilateral ischemic strokes affecting the middle cerebral artery (n=4, MCA), basal ganglia (n=2, BG), and brainstem (n=2, BS; Table 1). Three stroke participants were in the sub-acute phase (<6 months post-stroke), and the rest were chronic (>6 months). The numerical categories assigned to stroke participants reflect their motor deficit level, determined by the dynamic performance index; higher indices represent less severe deficits (Table 1).^4^

Participants were asked to reach in virtual reality (VR) shown in 3D environment created with Vizard (Worldviz) and shown via a head-mounted display (Oculus Rift), detailed in earlier publications (Fig. 1A).^4,12^ During reaching movements, motion capture data was recorded at 480 Hz and surface electromyography (EMG) was recorded at 2,000 Hz using MA400-28 (MotionLab Systems); both datasets were synchronized offline. EMG of twelve muscles was recorded including the clavicular head of pectoralis (Pec), teres major (TerM), anterior deltoid (ADel), posterior deltoid (PDel), the long and lateral heads of triceps (TriLo and TriLa), the short and long heads of biceps (BiS and BiL), brachioradialis (Br), flexor carpi radialis (FCR), flexor carpi ulnaris (FCU), and extensor carpi radialis (ECR, Fig. 1A). EMG data was analyzed in MATLAB (MathWorks Inc). The EMG data recorded from the Control group was reported by Korol and Gritsenko (2025).^27^ The kinematic and force data from the Stroke group was reported by Thomas et al. 2021.^4^

**Figure 1.**
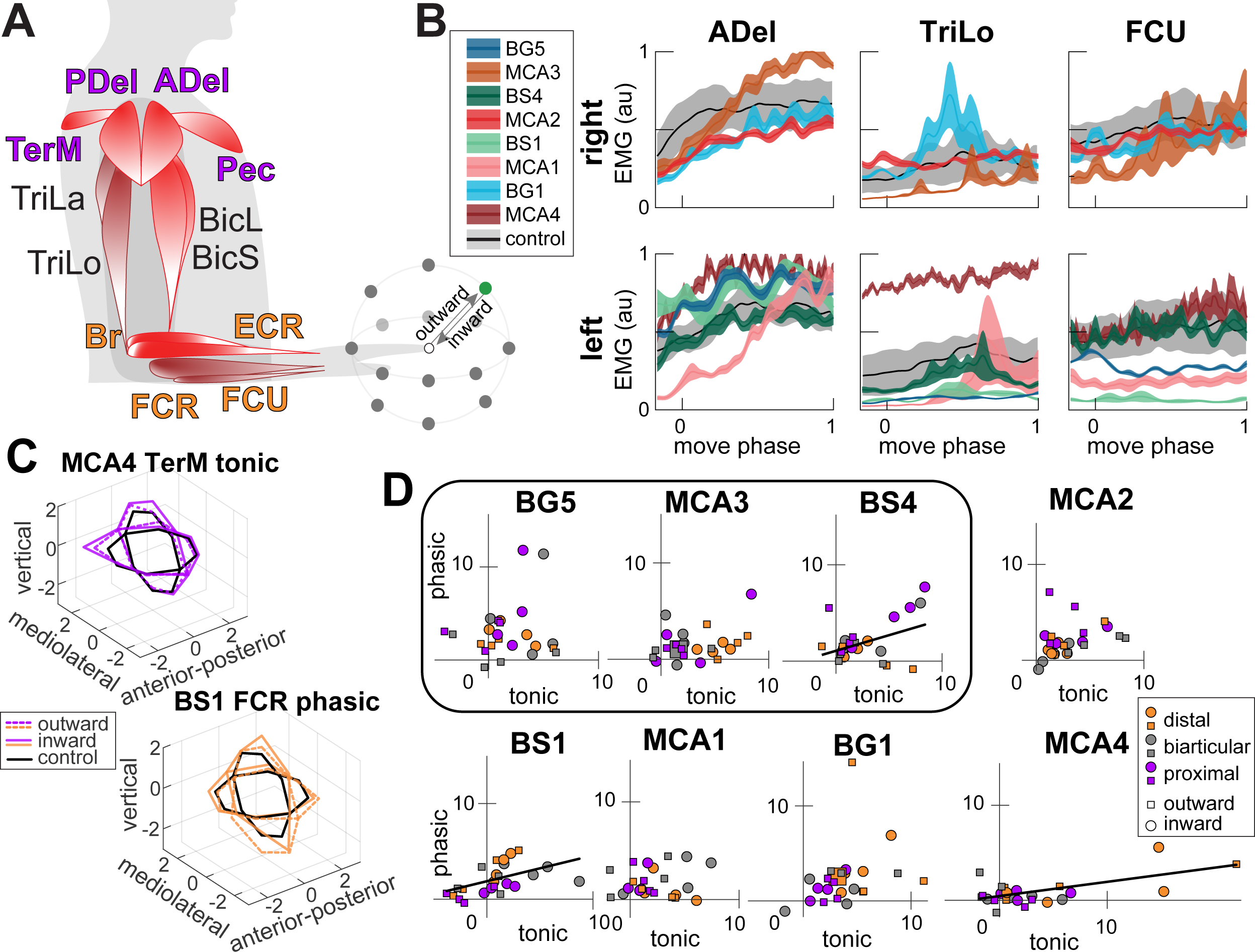
**A**. Experimental task side view. Each reaching movement was between the central target (open circle) and one of the peripheral targets (outward and inward reaches, green circle shows the exemplar reaching direction), target locations not to scale. Muscle activity was recorded from proximal muscles (purple labels): Pec, TerM, ADel, and PDel, biarticular muscles (black labels): TriLo, TriLa, BiS, and BiL, and distal muscles (orange labels): Brd, FCR, FCU, and ECR, abbreviations are as described in Methods. Bright red muscles support the arm against gravity in this task; darker-shaded muscles produce force in the direction of gravity. **B**. Examples of filtered and normalized muscle activity profiles for three muscles, abbreviated as in **A**, during outward reach to the green target in **A**. The gray shaded area shows the Control variance, and colors show the mean (line) and standard error (shaded area) for individual participants from the Stroke group. **C.** Plots show examples of tonic (top left) and phasic (bottom right) hemiparetic score differences (color lines) relative to controls (black lines) per reaching direction. The data were used to calculate the areas of the illustrated ellipses termed score tunings. **D.** Plots show tonic vs. phasic hemiparetic score tunings calculated from data shown in **C** per individual with stroke. The rounded rectangle outlines data for subacute participants. Squares indicate outward and circles indicate inward reaches; the color indicates values summed across proximal, biarticular, or distal muscles. Lines show significant (*p* < 0.05) linear regressions.

### Theory

The rotational motion of a joint with a single degree of freedom (DOF), driven by muscle forces, is governed by:

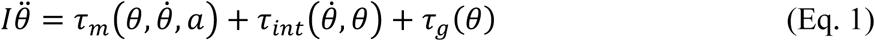

Here, *I* is segmental inertia; θ, 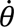, and 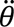 are joint angle, velocity, and acceleration; 𝜏_𝑚_ is the active muscle torque from neural activation 𝑎; 𝜏_g_ is gravitational torque and 𝜏_𝑖𝑛𝑡_includes passive torques that do not have a gravity-related component (e.g., interaction, Coriolis, and centrifugal moments). Only 𝜏_𝑚_ is directly controlled by the nervous system, which must compensate for body dynamics and gravity:

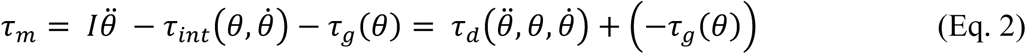

This decomposition separates the dynamic (𝜏_𝑑_) and anti-gravity (−𝜏_g_) components of the motor command.

Muscle torque about the *i*^th^ DOF is generated by the sum of forces from all *M* muscles crossing the joint, each scaled by their moment arm 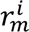:

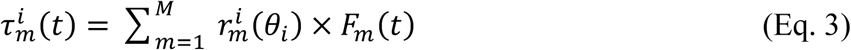

Here, the moment arms (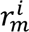) define agonist/antagonist actions and the relative contribution of each muscle’s force to the muscle torque. Muscle force 𝐹_𝑚_(t) is related to neural activation 𝑎_𝑚_(𝑡) through a Hill-type model:^28,29^

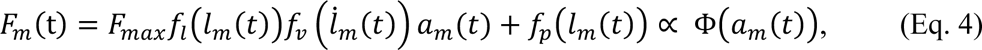

where 𝑓_𝑙_, 𝑓, and 𝑓_𝑝_ are nonlinear functions of muscle length 𝑙_𝑚_, velocity 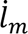, and passive properties, collapsed into Φ, a nonlinear mapping from activation to force.

When neural commands embed compensation for gravity (𝑎_𝑑_) and dynamics (𝑎_g_), muscle torques can be approximated as:

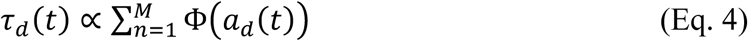

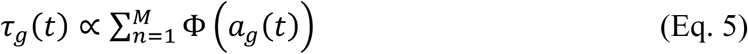

Thus, EMG-derived muscle recruitment can reflect biomechanical demands like gravity compensation and intersegmental coordination. We previously showed that tonic EMG aligns with 𝜏_g_, and phasic EMG with 𝜏_𝑑_.^12^ This analysis investigates how these components change after stroke.

### Data Analysis

EMG recordings were filtered (low-pass at 500 Hz, high-pass at 20 Hz, bandpass at 59-61 Hz), rectified, and further low-pass filtered at 10 Hz following SENIAM guidelines. EMG envelopes were normalized to movement duration, downsampled to 100 samples, and averaged across 15 repetitions per reaching direction for each muscle (total of 336 profiles). The sample size of 15 repetitions was derived empirically, it resulted in inter-trial variability in joint angle measurements of about 5 degrees.^30^ EMG amplitudes were scaled using maximum contraction values obtained across all reaching directions (Fig. 1B).

Principal component analysis (PCA) was used to separate tonic and phasic EMG components. Normalized, demeaned EMG profiles (12 muscles × 14 directions × inward/outward) were arranged into matrices (An×m; n = 100, m = 336). PCA assumptions for normality (Shapiro-Wilk test) and orthogonality (dot product calculation) were met, while homogeneity (Levene’s test) was not, indicating unequal variance across muscles and directions.

PCA utilized MATLAB’s singular value decomposition, extracting eigenvectors (principal components), variance accounted for (VAF), and scores. PCA was applied separately to data from Control and Stroke groups’ left and right arms (Control: left n=9, right n=9; Stroke: left paretic n=5, right paretic n=3, left non-paretic n=3, right non-paretic n=5). The PCA data is shared in a Figshare repository.^31^

Previous studies established that the first eigenvector represents tonic (gravity-related) EMG activity, while the second eigenvector captures phasic (intersegmental dynamic-related) EMG activity,^12^ both potentially altered by stroke. PCA scores quantify representation differences of EMG features across reaching directions and muscles. Due to minimal differences,^27^ Control scores from left and right arms were combined to enhance normative variability estimate. Paretic scores were then adjusted by subtracting bilateral control scores, measuring stroke-related deficits.

Score differences across reaching directions indicate feature recruitment variability dictated by task demands. The elliptical areas defined by these score differences, relative to controls, were calculated (Fig. 1C), providing quantitative measures of muscle over- or under-recruitment in stroke participants for tonic and phasic components.

### Statistics and Reproducibility

To determine the impact of stroke on the temporal evolution of the tonic and phasic biomechanical features in EMG, the first 2 eigenvectors and their VAF were compared between bilateral control data and the paretic and non-paretic data from Stroke group. We used the maximal coefficient of determination obtained from cross-correlation (*crosscorr* function in MATLAB Signal Processing Toolbox) between the mean control profile and the non-paretic or paretic individual profile for the 1^st^ and 2^nd^ eigenvector separately.

During reaching in certain directions, some muscles may have low tonic and phasic EMG, which would result in low PCA score values. Moreover, stroke can also cause reduced tonic and phasic EMG, which would also result in low scores. Here, we defined low scores as values below 5% of the maximal score. To evaluate the effect of stroke on score amplitude, we performed 2 two-sample t-tests comparing low scores of Control and Stroke groups for tonic and phasic components separately. The familywise error adjusted alpha was equal to 0.0253 using Bonferroni-Sidak correction.

We hypothesized that 1) stroke affects the recruitment of distal muscles that move the wrist and hand more than proximal and biarticular muscles that move the shoulder and elbow; 2) stroke affects the phasic component of EMG more than the tonic component. To test these hypotheses, we fitted a non-parametric generalized linear mixed-effects (GLME) model (*fitglme* function in Statistics and Machine Learning Toolbox, MATLAB) with an identity link and normal error distribution to examine the effects of three continuous predictors—Component (Tonic vs. Phasic), Direction (Outward vs. Inward), and Muscle Group (Proximal, Biarticular, vs. Distal)—on the sum of score tunings, along with their interactions. One-sample Kolmogorov-Smirnov test was performed on the GLME model residuals to check that the normality assumption is not violated (alpha = 0.05; *p* = 0.58). Random effects variance components were estimated for the intercept across subjects and for the residual error. The table below shows beta coefficients (β) that represent the estimated effects of predictors (fixed effects) on the outcome variable, adjusted for the random effects structure in the model and the associated standard error (SE), confidence intervals (CI), and p-values. The hypotheses will be rejected if the beta values are insignificant at alpha = 0.05.

To test if stroke affects both tonic and phasic EMG in individual participants, we regressed tonic vs. phasic score tunings using linear regression analysis (*regress* function in MATLAB Statistics and Machine Learning Toolbox). We excluded outliers in a percentile threshold from 2.5% to 97.5% (*isoutlier* function in MATLAB Data Preprocessing Toolbox).

To evaluate how stroke deficits measured from joint moments in our earlier study are related to the EMG deficits, we regressed the score tunings for tonic and phasic components during outward and inward reaches against gravitational and dynamic torque performance index. We have also regressed the score tunings against the years since the stroke.

The hemiparetic score values change across reaching directions in proportion to how much a given muscle is over- or under-recruited for a given movement. To determine whether these abnormal recruitment patterns are similar across muscles, we regressed the hemiparetic score differences for 28 reaching directions between each muscle pair for tonic and phasic EMG components separately. We excluded outliers that is more than three median absolute deviations from the median (*isoutlier* function in MATLAB Data Preprocessing Toolbox). This was achieved with linear regression analysis for Control and paretic arm Stroke Groups with adjusted alpha = 0.0008 (Bonferroni-Sidak correction). Linear regression analysis was performed using *regress* function in MATLAB Statistics and Machine Learning Toolbox. Each matrix, ***C_nxm_***, consisted of n = 28 reaching directions by m = 12 muscles per participant for tonic and phasic EMG components. Significant relationships were indicated by moderate and high coefficients of determination (R^2^) for each muscle pair. The resulting matrix of R^2^ values, represented as heatmaps in Figures 3 and 4, show how many muscles were over- or under-recruited together in each participant. We also preserved the sign of the regression slope to indicate whether the coupled recruitment represents co-contraction (positive sign) or antagonistic activation (negative sign) of the muscle pair.

Sensitivity analysis: to evaluate the effect of differences between dominant and non-dominant motor control on the estimate of stroke deficits, the same analyses were applied to hemiparetic scores normalized to matched data from left- or -right arm of control participants. The results were the same.

## Results

We recruited 19 potentially eligible participants, 2 of whom were ineligible due to trauma to the arm (n=1) and the absence of stroke diagnosis in the medical records (n=1). The remaining 17 eligible participants all completed the study and showed consistent motion and joint moment profiles^4,12^ but more variable surface EMG profiles (Fig. 1B).^27^

To understand how muscle recruitment changes after a stroke, we compared the features identified by PCA in individuals in the Control and Stroke groups. Our first step was to confirm that the principal components (i.e., eigenvectors) derived from EMG data in both groups are oriented in similar directions. We found that the temporal patterns of both primary features of the Stroke group were similar to the corresponding features in the Control group (Supplemental Fig. S1A), and they captured a similar amount of variance in EMG (Supplemental Fig. S1B). The variance accounted for (VAF) by the first two principal components from EMG of paretic muscles in the Stroke group was equal to 53% ± 13% and 17% ± 5% (mean ± standard deviation across participants) for 1^st^ (tonic) and 2^nd^ (phasic), respectively. The corresponding values for the Control group were 52% ± 7% and 16% ± 5%, respectively, and for the non-paretic arm in the Stroke group were 54% ± 10% and 19% ± 7%, respectively. This implies that the underlying data structure across individuals shares a common pattern of variance. This agrees with subjective observations of clinicians and scientists that the patterns of post-stroke EMG often appear normal despite motor deficits.

Next, we analyzed the principal component scores, where the amplitude reflects how strongly a specific biomechanical feature — related to gravity or intersegmental dynamics — is expressed in a muscle’s EMG activity during movement. Low scores indicate that the feature accounts for little variance in the EMG signal for certain muscles and reaching directions, suggesting abnormal recruitment patterns in the hemiparetic arm after stroke. Our data show that, in control participants, 18% ± 7% (mean ± SD across participants) of tonic scores were low, or below 5% of the maximum, compared to 23% ± 9% for the paretic arm of participants with stroke. A two-sample t-test showed no significant difference between groups in the proportion of low tonic scores, t(24) = −1.31, *p* = 0.2013. However, the percentage of low tonic hemiparetic scores across individuals increased with time since stroke (Pearson correlation coefficient R = 0.85, *p* = 0.0068). Moreover, a significantly higher proportion of phasic hemiparetic scores were low (30% ± 6%) compared to controls (24% ± 6%) (two-sample t-test, t(24) = 2.69, *p* = 0.0127). There was no linear relationship between the percentage of low phasic hemiparetic scores and time since stroke (R = 0.01, *p* = 0.9812). This suggests that stroke reduces both components of EMG activity, though to varying degrees depending on individual pathophysiology; the phasic component associated with intersegmental dynamics is most affected.

Altered recruitment patterns were also evident across different reaching directions, each requiring varying combinations of forces for arm support against gravity and for reaching toward or away from a target. Consequently, the expression of biomechanical features in a given muscle’s EMG signal varied with reaching direction—a directional variability that was clearly observable in hemiparetic score differences after stroke (Supplementary Figs. S2, S3). We captured these differences in muscle recruitment as score tunings (see Methods above). In this analysis, negative tunings indicate a reduced representation of the feature in the hemiparetic EMG or under-recruitment of muscles, while positive tunings indicate an increased representation or over-recruitment of muscles relative to controls. The resulting score tunings were mainly positive with individual variability in the extend of deficit in tonic and phasic recruitment of proximal, biarticular, and distal muscle groups (Fig. 1D). In three participants (BS4, BS1, and MCA4), the phasic and tonic score tunings were correlated. The variation in tonic and phasic recruitment deficits across individuals indicates that stroke can impair either mechanism independently or both simultaneously. The GLME model showed a significant intercept (β = 3.85, p < 0.001), indicating a nonzero baseline value for the muscle recruitment deficits captured by tonic and phasic score tunings. The Component predictor had a significant negative effect (β = −0.54, p = 0.020), showing that the phasic EMG scores (2^nd^ principal component) were not surprisingly lower than the tonic EMG scores (1^st^ principal component). The Muscle Group predictor also had a significant negative effect (β = −0.48, p = 0.005), meaning distal and biarticular muscles had lower deficits than proximal muscles. The interaction Component × Muscle Group was significant and positive (β = 0.24, p = 0.027), implying that the negative effect of Component depends on the level of Muscle Group — the differences between phasic and tonic recruitment is lower in biarticular and distal muscles compared to proximal muscles. Other predictors and interactions were not statistically significant (Table 2). Random effects analysis showed small variability across participants (SD = 0.09), and residual error SD was 0.42. This shows that after stroke, the damaged motor system over-activates many muscles in the hemiparetic arm—particularly for the action to support the limb against gravity— making movements energy-inefficient and likely contributing to fatigue.

Deficits in tonic muscle recruitment varied with the time since stroke. However, the deficits in tonic recruitment of proximal muscles were markedly different for outward and inward movements (Fig. 2A), indicating different neural control mechanisms for outward reaching and returning retrieval movements. In chronic stroke participants, tonic overactivation of hemiparetic biarticular muscles during outward reaches decreased as the number of years since stroke increased (*R^2^* = 0.90, *p* = 0.019; Fig. 2A). This trend was especially evident in the short head of triceps (*R^2^* = 0.30, *p* = 0.027). Furthermore, the difference in tonic activation between proximal and distal muscles increased with time post-stroke (*R^2^* = 0.45, *p* = 0.019) with differences between outward and inward reaches diminishing (Fig. 2C). This was driven by increasing tonic overactivation of proximal muscles and decreasing tonic overactivation of distal muscles, but each of these regressions with years since stroke were not significant (chronic proximal: *R^2^* = 0.16, *p* =0.245; chronic distal: *R^2^* = 0.35, *p* = 0.070). In subacute participants, tonic muscle activation showed marked differences between outward and inward reaches. Outward reaches were dominated by proximal over-recruitment, with summed proximal muscle score tunings of 6.70, 11.56, and 25.09 and corresponding distal muscle differences of – 2.4, 2.90, and 6.04 for subjects BG5, BS4, and MCA3, respectively. In contrast, inward reaches exhibited more variable deficits in proximal and distal muscle recruitment, with summed score tunings of 9.29, 6.69, and 20.72 for proximal muscles, and 9.59, 22.66, and 12.36 for distal muscles in the same subacute individuals. Taken together, the data support clinical observations that subacute stroke phase is distinct from the chronic phase. As more time passes after stroke, the tonic over-recruitment of proximal muscle tends to increase or stay the same while the tonic over-recruitment of biarticular and distal muscles tends to decline.

**Figure 2.**
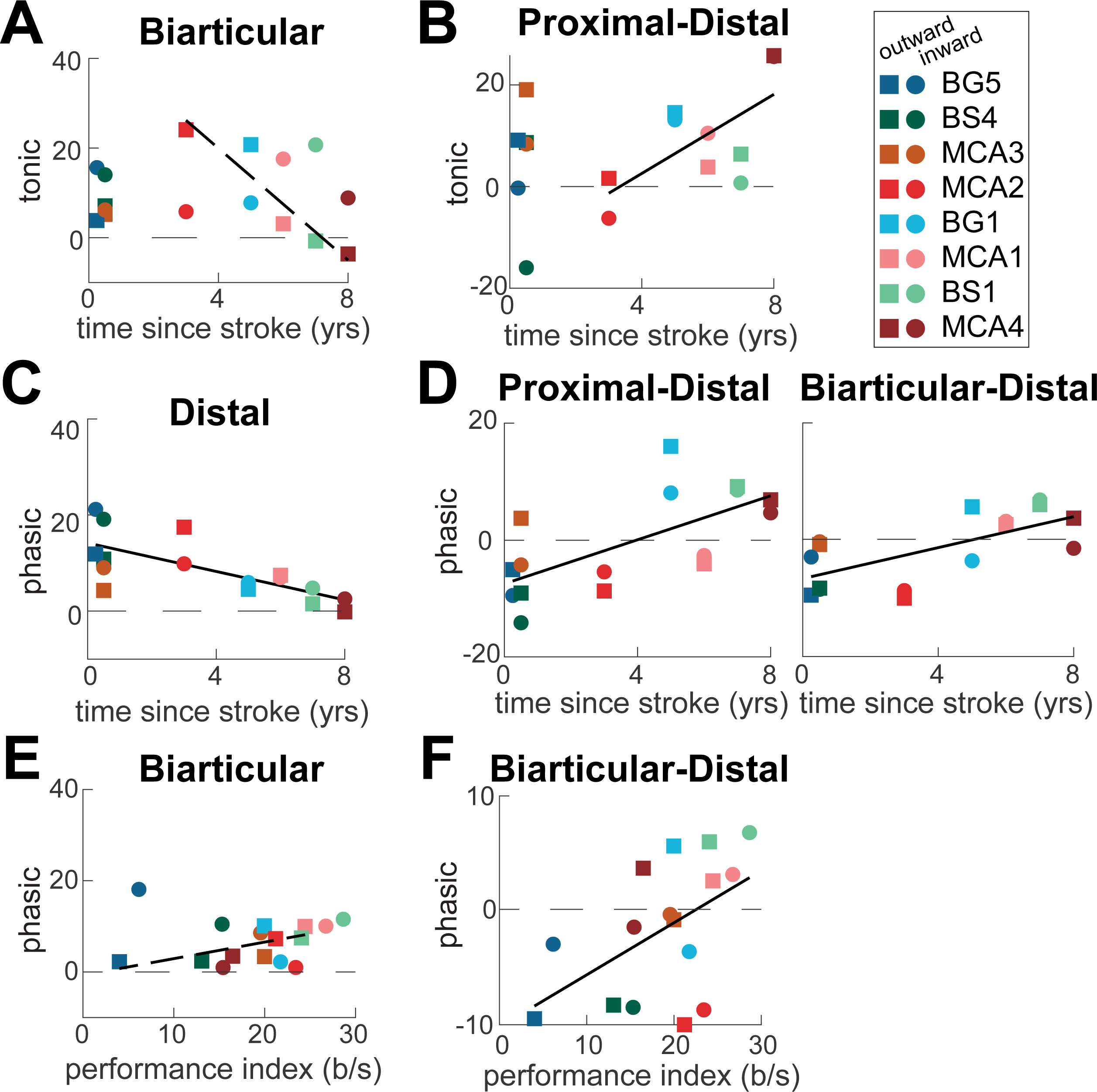
Deficits in muscle recruitment. **A.** Plot shows tonic score tunings for the biarticular muscles plotted against the number of years post-stroke. Squares indicate outward and circles indicate inward reaches for individual participants; color coded as in Fig. 1. The dashed line shows a significant (*p* < 0.05) linear regression for the outward reaches by chronic participants. **B.** Plot shows tonic score tunings calculated between proximal and distal muscles plotted against the number of years post-stroke, styled as in **A**. The line shows a significant linear regression across all reaching directions by chronic participants. **C.** Plot shows phasic score tunings for the distal muscles plotted against the number of years post-stroke, styled as in **A**. The line shows a significant linear regression across all reaching directions and participants. **D.** Plots show phasic score tunings calculated between proximal and distal (left plot) and biarticular and distal (right plot) muscles plotted against the number of years post-stroke, styled as in **A**. Lines show significant linear regressions across all reaching directions and participants. **E.** Plot shows phasic hemiparetic score tunings for the biarticular muscles plotted against the dynamic performance index, styled as in **A**. The dashed line shows a significant linear regression for the outward reaches by chronic participants. **F.** Plot shows phasic hemiparetic score tunings calculated between biarticular and distal muscles plotted against the dynamic performance index, styled as in **A**. The line shows a significant linear regression across all reaching directions and participants.

Deficits in phasic muscle recruitment also varied with the time since stroke. The phasic over-activation of distal muscles decreased across all participants (*R^2^* = 0.52, *p* = 0.002; Fig. 2B). These trends were present in flexor carpi radialis (*R^2^* = 0.40, *p* = 0.009) and flexor carpi ulnaris (*R^2^* = 0.36, *p* = 0.015). Moreover, the difference in phasic over-activation between proximal and distal muscles (*R^2^* = 0.44, *p* = 0.005) and between biarticular and distal muscles (*R^2^* = 0.46, *p* = 0.004) also increased with time post-stroke (Fig. 2D). These trends were primarily driven by the progressive decrease in phasic over-recruitment of distal muscles in individuals further removed from the time of stroke onset, consistent with the findings in Fig. 2B. Moreover, the differences between proximal and distal muscles in phasic and tonic activation were correlated across all participants (*R^2^* = 0.38, *p* = 0.011), indicating a similar underlying neural mechanism of decreasing both tonic and phasic over-recruitment of distal muscles. Overall, these results suggest that adaptation over time since stroke is most evident in the reduced activation of distal muscles compare to the activation of proximal and biarticular muscles, especially in the phasic component of muscle recruitment.

Stroke-induced motor deficits during reaching are reflected in muscle torques, which represent the forces required to move the limb. These torques can be further decomposed into components corresponding to the forces necessary for arm support against gravity and those required to propel the limb during reaching. Given that muscle torques arise from the combined actions of all muscle groups, our EMG analysis enabled quantification of the contributions of specific muscle groups to post-stroke motor impairments. We have shown in an earlier study that the performance index derived from the dynamic component of muscle torques, which is similar to the phasic component of EMG,^12^ is a sensitive indicator of the severity of post-stroke motor deficits.^4^ Here too we have found that the severity of motor deficits measured with the performance index derived from muscle torques was linearly related to the number of years post-stroke (gravitational performance index vs. years: *R^2^* = 0.27, *p* =0.039; dynamic performance index vs. years: *R^2^* =0.32, *p* = 0.023). Moreover, the phasic over-recruitment of biarticular muscles during outward reaches exhibited a strong linear relationship with the summed performance index derived from the dynamic component of shoulder and elbow muscle torques (*R^2^* = 0.55, *p* = 0.036; Fig. 2E). This association was particularly notable in the short head of the biceps (*R^2^* = 0.28, *p* = 0.034) and showed an inverse relationship in the flexor carpi radialis (*R^2^* = 0.29, *p* = 0.031) across both outward and inward reaches. Furthermore, phasic score differences between biarticular and distal muscles were linearly related to the dynamic performance index (*R^2^* = 0.27, *p* = 0.038; Fig. 2F). Collectively, these findings demonstrate that stroke-related changes in muscle recruitment patterns directly contribute to impairments in generating the forces necessary for limb propulsion and coordinating movement across joints during reaching tasks.

The results above show that post-stroke muscle activation deficits affect entire muscle groups rather than individual muscles. To explore this further, we examined muscle coactivation—that is, the simultaneous recruitment of multiple muscles to perform a given task. The coactivation between antagonistic muscles also contributes to joint or limb stiffness, an important parameter controlled by the central nervous system. In previous work, we showed that in control participants, tonic activity in biarticular and distal muscles, especially in the left arm, was broadly correlated across reaching directions, reflecting their coactivation to support the arm against gravity.^27^ However, there was substantial variability across individuals, with two control participants showing few co-activating muscle pairs in the left arm (Fig. 3A). After stroke, the number of tonically co-activating muscle pairs in left-paretic participants was near the lower end of the control range, whereas in right-paretic participants, it was near the upper end (Fig. 3B, C). In contrast, control participants showed minimal phasic coactivation in both arms^27^ (Fig. 4A), and this pattern remained largely unchanged after stroke, with few phasically co-activating muscle pairs observed (Fig. 4B, C). Notably, we found a significant inverse linear relationship between the number of phasically co-activating proximal and distal muscles and the dynamic performance index (*R^2^* = 0.67, *p* = 0.013), indicating that greater impairments in generating propulsive forces are associated with more abnormal coactivation between proximal and distal muscles.

**Figure 3.**
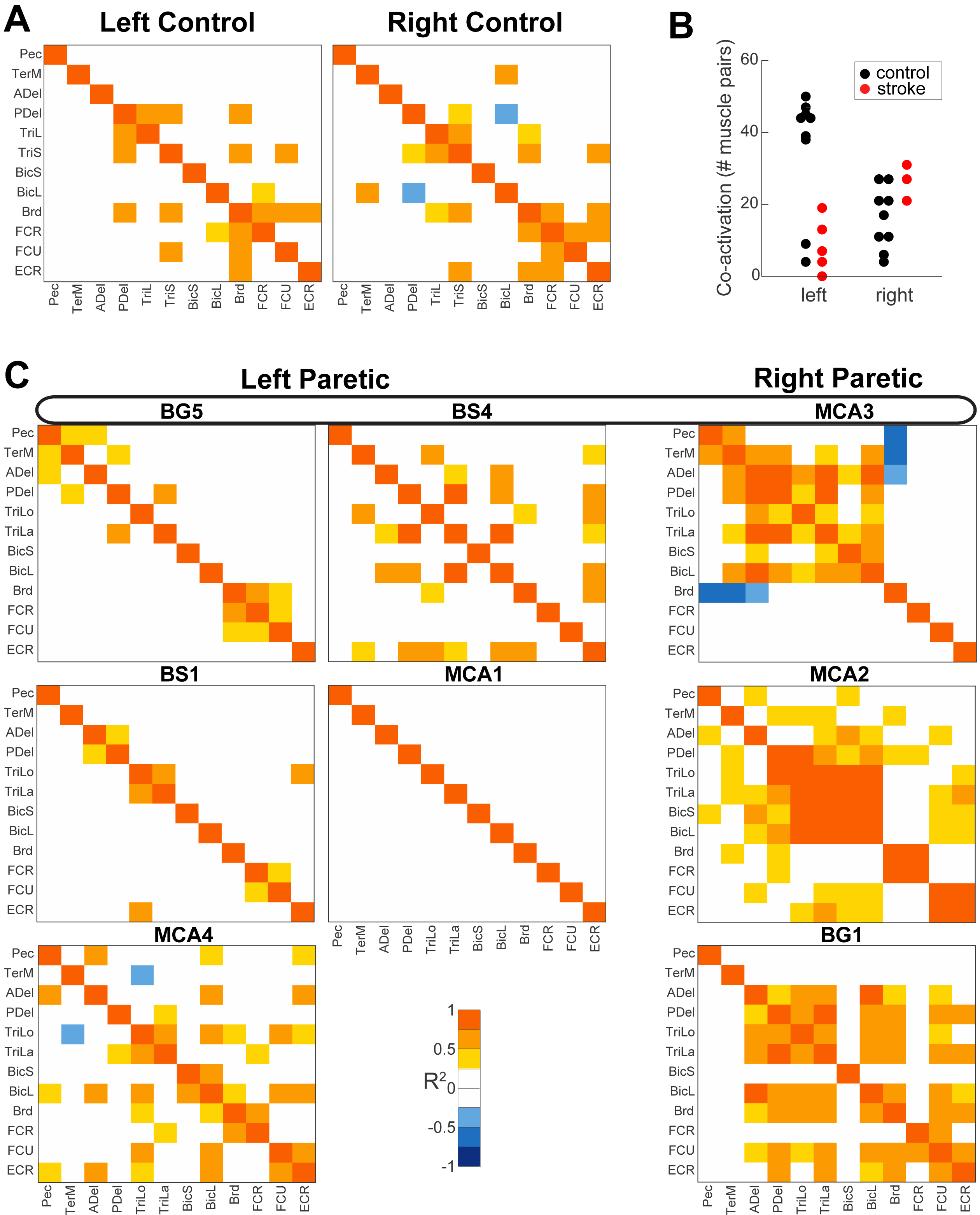
Tonic coactivation. **A.** Heatmaps show matrices of significant regressions between tonic scores of muscle pairs across reaching directions with the left (left plot) and right (right plot) limb by a young control participant (S5 from Korol et al. 2025) who showed fewer coactivations between muscles of their left arm. The colors show the values of the coefficient of determination (R^2^) for regressions between the tonic scores of muscle pairs across reaches in multiple directions. **B.** Symbols show the number of moderate and strong (R^2^ >0.5) muscle coactivations in individuals with (red) and without (black) stroke. **C.** Heatmaps of tonic muscle coactivations for participants with stroke during reaching with their paretic arm. Colors are as in **A**.

**Figure 4.**
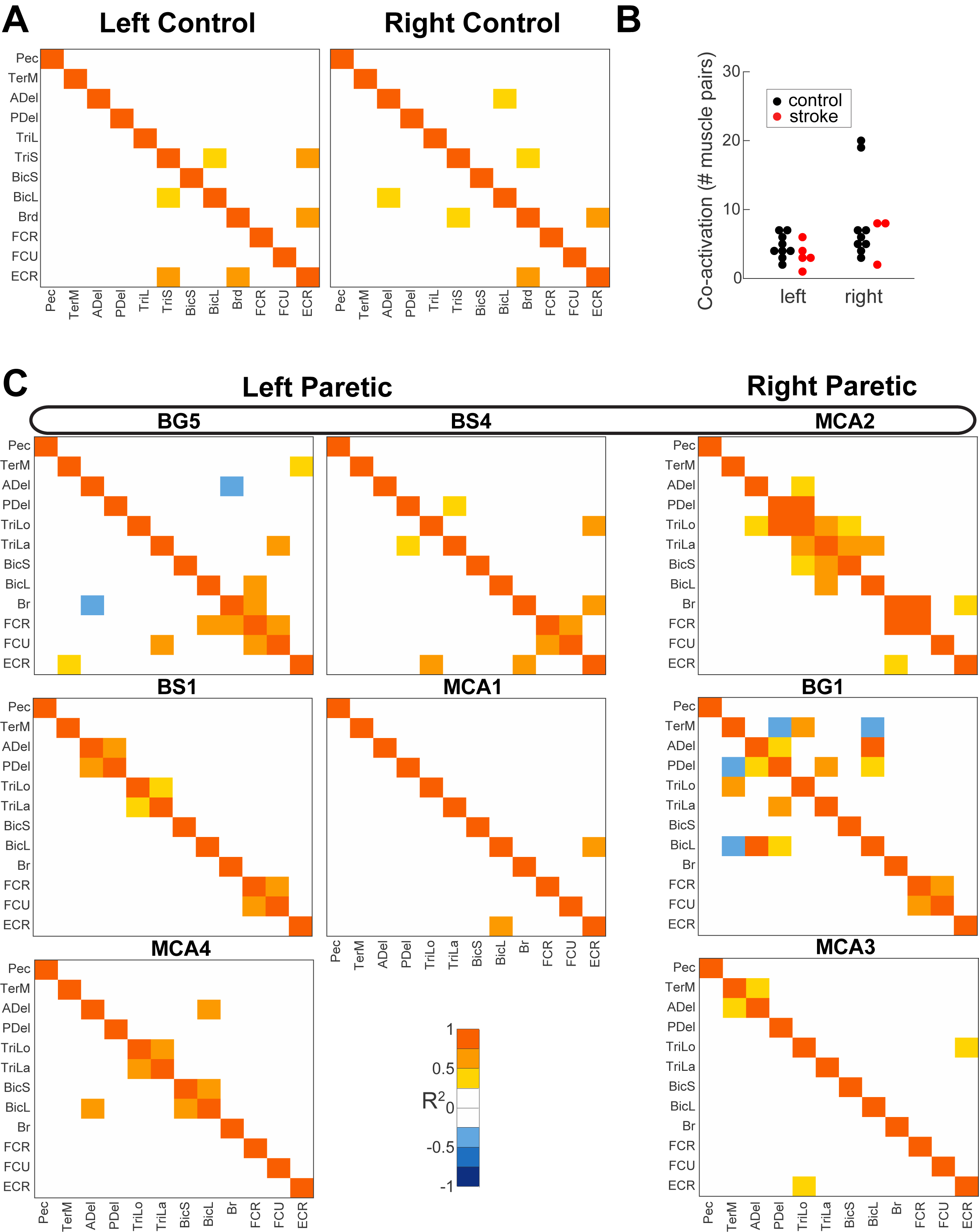
Phasic coactivation. **A.** Heatmaps show matrices of significant regressions between phasic scores of muscle pairs across reaching directions with the left (left plot) and right (right plot) limb by a young control participant (S5 from Korol et al. 2025). The colors show the value of the coefficient of determination (R^2^) for regressions between the phasic scores of muscle pairs across reaches in multiple directions. **B.** Symbols show the number of moderate and strong (R^2^ >0.5) muscle coactivations in individuals with (red) and without (black) stroke. **C.** Heatmaps of phasic muscle coactivations for participants with stroke during reaching with their paretic arm. Colors are as in **A**.

## Discussion

Our results show that stroke disrupts muscle recruitment during reaching, even among individuals with mild impairments. Distal muscle impairment severity increases over time post-stroke, but proximal muscles are also abnormally over-recruited during movements that usually require less activation, indicating disrupted directional tuning. Additionally, paretic muscle coactivation between proximal and distal muscles is abnormal. These findings reject the hypothesis that stroke primarily impacts distal muscles more severely than proximal or biarticular muscles, as both groups are variably affected.

Our results further reject the hypothesis that stroke disproportionately affects the phasic component of EMG compared to the tonic component. Instead, deficits in phasic and tonic recruitment correlate linearly in some participants, and phasic and tonic differences between proximal and distal recruitment correlate across participants and align with overall reaching impairments. This suggests that stroke prompts widespread muscle over-activation, particularly for supporting the limb against gravity, resulting in inefficient movements and potential fatigue.

Our data clarify how stroke damages specific neural control pathways, influencing motor impairment and recovery. Our results suggest that the natural organization of muscles into proximal and distal groups,^32^ is differentially impacted by stroke. This may be caused by the individual stroke-related damage to the reticulospinal pathways that contributes to abnormal flexor synergies and over-engagement of proximal muscles^17,33^ and to the corticospinal pathway that governs distal fine motor control.^34^ The limitation of this conclusion is the small sample size of the study. Power analysis (MATLAB function *sampsizepwr*) indicates that to detect the mean change of 0.478 ± 1.027 SD in post-stroke recruitment between muscle groups (Table 1) with 80% power, the study needs 30 participants with stroke. Future studies with this sample sizes will reveal the details of how the plasticity in the two pathways affect muscle recruitment after stroke.

Identifying underlying causes of movement inefficiency and fatigue remains challenging, but assessing mechanical impedance provides a quantitative framework.^35^ Impedance, comprising stiffness and viscosity, increases during unstable tasks or when precision is required.^36–38^ Our results show that after stroke, lateralization of impedance control is disrupted: right-hemisphere strokes reduce consistent muscle coactivations typical of non-dominant hemisphere-driven gravity compensation,^27^ while left-hemisphere strokes increase coactivation. Greater deficits in generating propulsive forces also correlate with abnormal coactivation patterns, underscoring that stroke-induced disruption in impedance control significantly contributes to motor inefficiency and fatigue. Quantifying such disruptions may help pinpoint their underlying neural causes and guide more targeted rehabilitation strategies.

## Author Contributions

ASK AKA VG

## Statements and Declarations

The study protocol was approved by West Virginia University’s Institutional Review Board (Protocol #1311129283). Participants were recruited from Morgantown, WV area between 2014 and 2017. All participants came provided written informed consent.

## Consent for Publication

Not applicable.

## Data availability

The data is available on Figshare DOI:

