## Supplemental for "Stroke-Related Changes in Tonic and Phasic Muscle Recruitment During Reaching Reveal Pathway-Specific Motor Deficits"

### Supplemental Figures

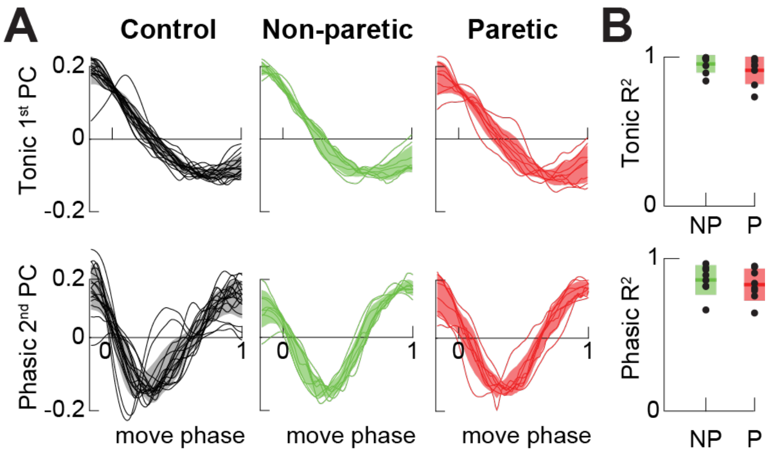


**Supplemental Figure S1. A**. Principal components' temporal profiles, where the profile of the Tonic 1st principal component (PC) related to gravity compensation is plotted in the top row and the profile of the Phasic 2nd PC related to intersegmental dynamics is plotted in the bottom row. Solid lines represent each participant. The shaded area is the standard deviation across participants. Black is a control arm (right and left), green is the non-paretic arm, and red is the paretic arm. **B**. The coefficient of determination ($R^2$) between corresponding profiles in **A**; NP stands for non-paretic and P stands for paretic.

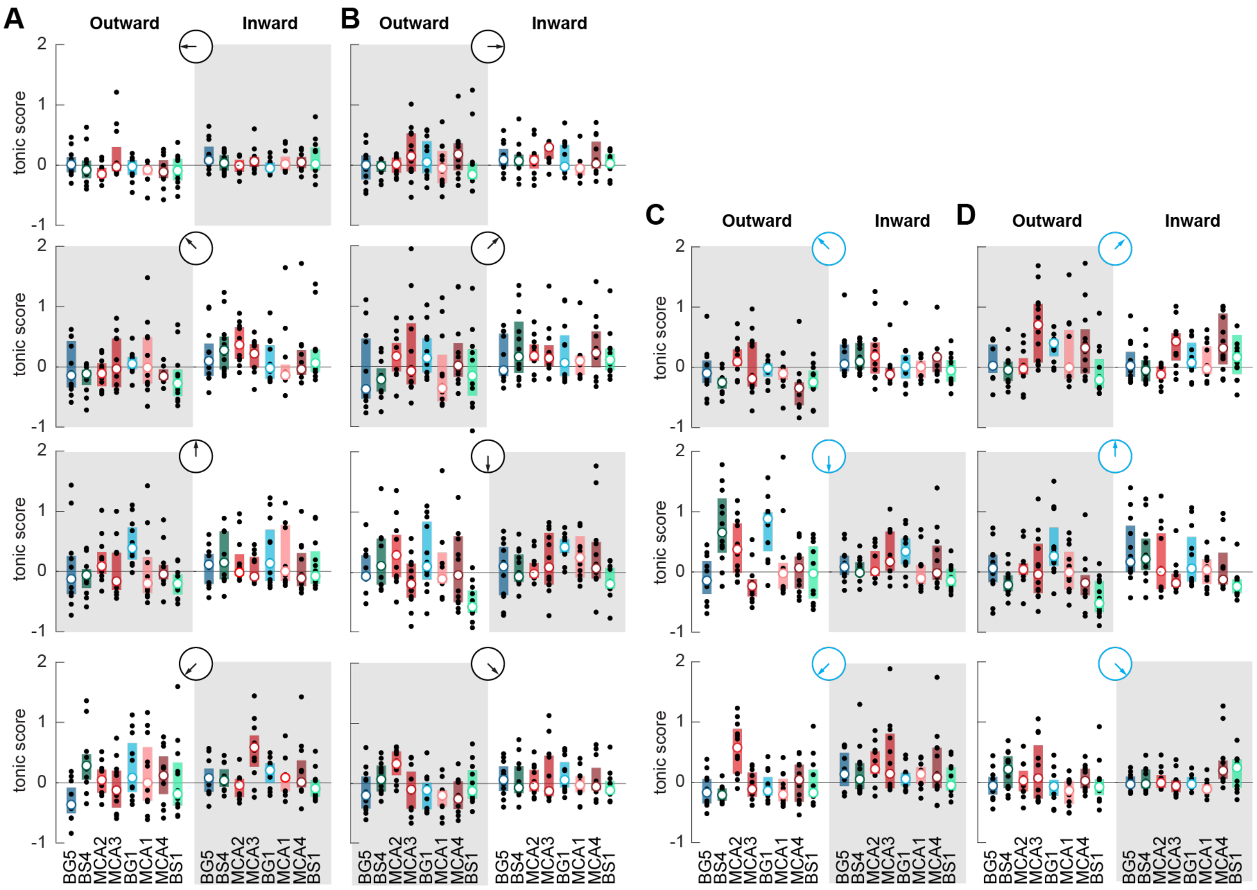


**Figure S2. Post-stroke tonic score differences per reaching direction.** Black dots show differences between each participant with stroke and corresponding bilateral data from control participants averaged across movement directions. Open symbols are medians, and shaded boxes show 25% and 75% interquartile ranges across participants with stroke. Coded IDs of participants with stroke are arranged by the months/years after a stroke. Gray shaded areas highlight movement directions against gravity. **A.** Plots show data for movements to targets on the left side of horizonal transverse plane; pictograms show view from above with participant located on the left of the pictogram. **B.** Plots show data for movements opposite to those in **A. C.** Plots show data for movements to targets on the left side of vertical sagittal plane, pictograms show view from side with participant located on the left of the pictogram. **D**. Plots show data for movements opposite to those in **C.**

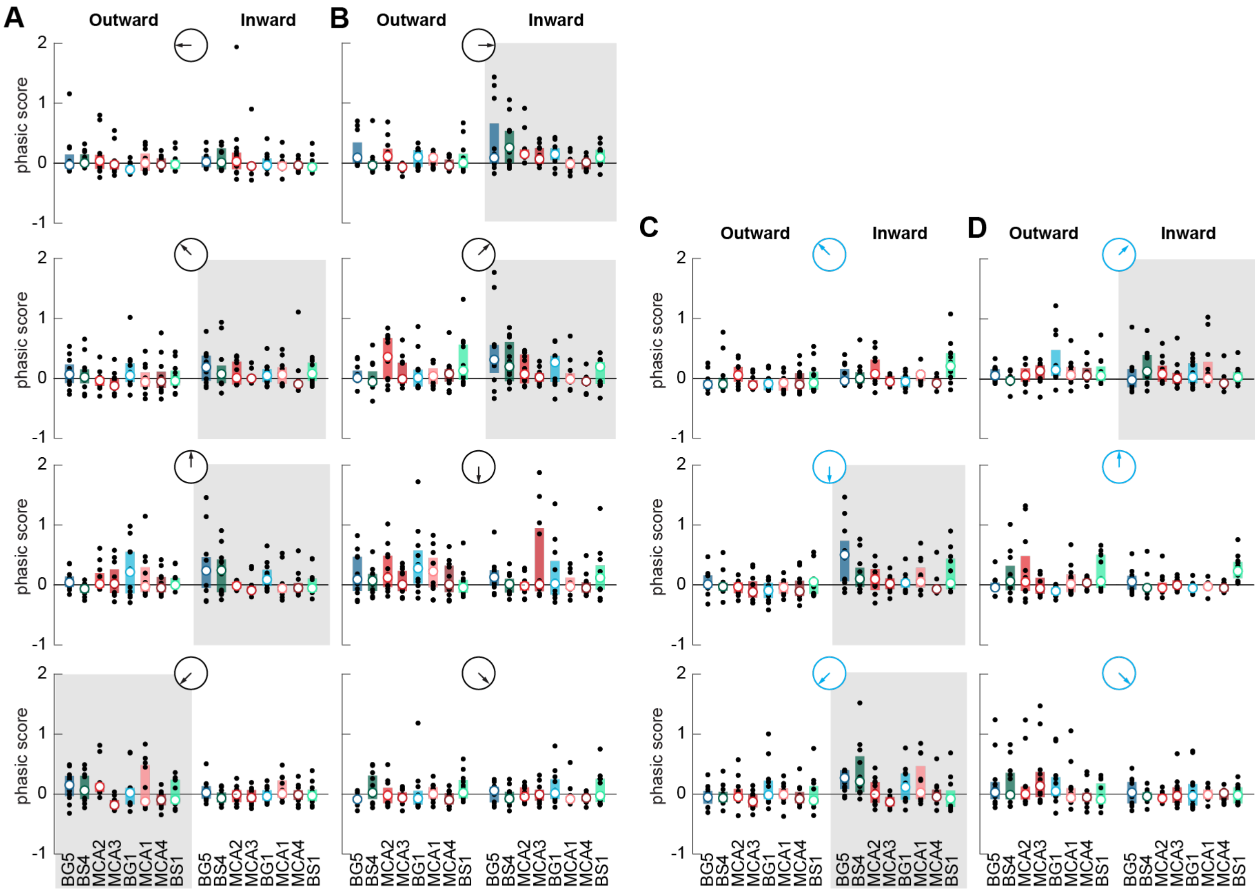


**Figure S3. Post-stroke phasic score differences per reaching direction.** Black dots show differences between each participant with stroke and corresponding bilateral data from control participants averaged across movement directions. Open symbols are medians, and shaded boxes show 25% and 75% interquartile ranges across participants with stroke. Coded IDs of participants with stroke are arranged by the months/years after a stroke. Gray shaded boxes highlight movement directions that show decreasing scores with time after stroke. **A.** Plots show data for movements to targets on the left side of horizonal transverse plane, pictograms show view from above with participant located on the left of the pictogram. **B.** Plots show data for movements opposite to those in **A. C.** Plots show data for movements to targets on the left side of vertical sagittal plane; pictograms show view from side with participant located on the left of the pictogram. **D.** Plots show data for movements opposite to those in **C.**
